# The High-resolution Timeline of Expression of Ribosomal Protein Genes in Yeast

**DOI:** 10.1101/170399

**Authors:** Xueling Li, Gang Chen, Bernard Fongang, Dirar Homouz, Maga Rowicka, Andrzej Kudlicki

**Affiliations:** Department of Biochemistry and Molecular Biology, University of Texas Medical Branch, Galveston, TX 77555, USA; Institute for Translational Sciences, University of Texas Medical Branch, Galveston, TX 77555, USA; Hefei Institute of Intelligent Machines, Institutes of Physical Science, Chinese Academy of Sciences, Hefei, China.; Department of Computer Science, School of Information Science and Engineering, Central South University, Changsha, China; Khalifa University, Abu Dhabi, UAE; Sealy Center for Molecular Medicine, University of Texas Medical Branch, Galveston, TX 77555, USA

## Abstract

The yeast ribosome is a complex molecular machine built from four rRNAs and over 70 r-proteins. Ribosome biogenesis involves ordered incorporation of ribosomal proteins, accompanied by and association and dissociation of other proteins specific to different stages of the process. By model-based analysis of temporal profiles of gene expression in a metabolically regulated system, we obtained an accurate, high-resolution estimation of the time of expression of genes coding for proteins involved in ribosome biogenesis. The ribosomal proteins are expressed in a sequence that spans approximately 25-minutes under metabolically regulated conditions. The genes coding for proteins incorporated into the mature ribosome are expressed significantly later than those that are not incorporated, but are otherwise involved in ribosome biogenesis, localization and assembly, rRNA processing and translational initiation. The relative expression time of proteins localized within specified neighborhood is significantly correlated with the distance to the centroid of the mature ribosome: protein localized closer to the center of mass of the entire complex tend to be expressed earlier than the protein localized further from the center. The timeline of gene expression also agrees with the known dependencies between recruitment of specific proteins into the mature ribosome. These findings are consistent in two independent experiments. We have further identified regulatory elements correlated with the time of regulation, including a possible dependence of expression time on the position of the RAP1 binding site within the 5’UTR.

## Introduction

The eukaryotic ribosome is an ancient and conserved protein synthesis machine that translates messenger RNAs into proteins. The yeast ribosome consists of four ribosomal RNAs (rRNAs) and 79 proteins (r-proteins) [1]. The biogenesis and assembly of a ribosome requires 76 different small nucleolar RNAs, more than 200 assembly factors, in addition to the r-ribosomal proteins [2]. This regulation of ribosome biogenesis is largely transcriptional. Several transcription factors and regulatory motifs are involved in the regulation of r-protein expression, including: Rap1/SFP1[3,4,5], A-rich [6], IFH1/FHL1 [7], Motif213 associated with Rap1 [8], AAAAATTTT [9], and GATGAG [9,10].

It has been shown that the maturation of molecular complexes may be accompanied by just-in-time expression of their building block proteins, regulated at the transcript level [11,12,13]. According to this paradigm, the protein components of a macromolecular complex are generally expressed precisely when they are needed. We have developed and successfully utilized model-based algorithms for determining the time of expression peak with high resolution and accuracy from sparsely sampled time course expression data [11,14]. We have demonstrated that the relative times of expression for components in transcriptional, regulatory, and signaling pathways [11,15], may correspond to the causal relations between genes. We also noticed that expression levels of ribosomal protein genes are strongly modulated during the yeast metabolic cycle (YMC). The expression profiles of ribosomal genes measured in [11] have been reproduced in a recent RNA-Seq based study of the YMC [16]. We therefore expect that the just-in-time expression paradigm may also apply to the genes encoding r-proteins incorporated into the mature yeast ribosome.

Ribosomes are produced in a three-stage process, comprising the assembly of 90S large pre-preribosome particles containing mostly pre-40S subunit related components and a few 60S subunit components in the nucleolus, pre-60S and pre-40S pre-ribosome particles in the nucleoplasm, and 80S mature ribosome in the cytoplasm [17,18,19]. In this work, we focus on characterizing the transcriptional timing of the r-proteins and associated proteins involved in ribosome biogenesis, and the correlation of the expression times with the functional and structural features of these proteins. The r-proteins in the ribosome show a hierarchical order of assembly [20], a process of assembly proceeding in a specified succession where the ribosome proteins from different neighborhoods are incorporated orderly. A correlation exists between r-protein locations and assembly order in both the small subunit (SSU) and large subunit (LSU) of mature ribosomes [21,22]. Since the structure of the ribosome is associated with the order of assembly, it may be expected that a correlation between relative positions of r-proteins within the structure and their timeline of expression will also be observed.

We previously demonstrated [11,15] that precise estimation of the time of a gene expression peak (timing of expression) is possible if the temporal profiles of the genes are produced by a process common for all the genes in question. In such cases, a model-based approach in gene expression time estimation will result in a resolution much higher than the resolution of the source data. This can be achieved because the method optimally uses the available prior information, and the timing is inferred from many independent measurements, reducing the impact of experimental errors. As a result, a correlation between the times of gene expression and the order of assembly was suggested in several macromolecular complexes [11]. The mRNA levels and expression times of may be tightly controlled during ribosome biogenesis. Their temporal expression profiles for most ribosomal genes are expected to be similar, however differences may exist that reflect the temporal order of expression. To investigate the expression timeline of ribosomal and ribosome-associated genes, we use the expression profiles during the yeast metabolic cycle (YMC). We chose this dataset because in this system the ribosome production is strongly and reproducibly regulated as a function of time [23,24]. Such strong modulation is not observed in other time course data like the cell cycle datasets [25,26,27,28,29], therefore expression profiles in cell-cycle synchronized cultures could not be used to infer timinig of ribosomal genes (See Methods and Supplementary Methods). During YMC, cell metabolism oscillates between an oxidative and a reductive phase [14,16]. The transition from the oxidative phase to reductive building phase is accompanied by substantial change in the proteome and is associated with a peak of increased ribosome production [14,30]. The temporal expression profiles of ribosomal genes during YMC are indeed similar [14,16], however differences in peak times between two functional r-protein gene groups are noted [16]. Here, we will precisely quantitate the differences in their times of expression that may uncover the logic underlying the timeline of expression of ribosomal genes. To this end, we first construct an empirical model of the temporal profile of expression of ribosomal proteins in YMC. By comparing this model with the expression profile of each gene, we estimated the expression times of ribosomal proteins and uncovered the precise timeline of the transcriptional program of ribosomal proteins.

Recent advances in structural biology have allowed crystallographic determination of structures of large macromolecular complexes, including bacterial [31], yeast [1], and human cytosolic ribosomes [32], as well as mitochondrial ribosomes of both human and yeast [33,34]. This made it possible to study the association between ribosome structure and its function and assembly [35]. To investigate whether the expression timeline of r-protein genes is associated with their ribosome assembly, we analyzed the timing results in the context of the crystallographic structure of the yeast 80S ribosome [1]. We studied the relationship between r-protein expression timing and the position of the mass center of the protein within the yeast 80S ribosome [1]. The rationale is that the r-proteins may have a spatial constraint for their neighboring r-protein, and that if the precise timeline of ribosomal protein gene expression is related to the spatial position of the corresponding r-proteins in the mature ribosome, the time of expression may serve a function in facilitating ribosome biogenesis and assembly. We compared the differences of expression times and differences of mass center distances of all defined neighboring pairs of ribosomal proteins and indeed found a significant correlation (see Results).

Furthermore, previous models of the molecular mechanisms controlling the time of expression use gene linear characteristics, such as the position and context of the cis-regulatory elements [8,12]. To explore the possible mechanism that regulates the precise timing of the r-protein expression, we searched for regulatory elements whose presence or position is correlated with time expression; such elements could potentially be involved in controlling a precisely timed sequence of gene expression.

Finally, to validate the reproducibility of estimated high-resolution precise timing, we further examined the correlation between the estimated times from YMC data based on different platforms and sampling, i.e., the microarray [11] used here and the recent RNA-Seq YMC [16].

## Results

### Model-based timing of expression peaks

We have implemented an algorithm for inferring the precise times of gene expression (see Methods). Our approach is based on the assumption that the genes have similar temporal expression profiles, with individual time-shifts 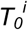. The gene-specific profile for gene (*i*) will then read 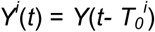, where *Y*(·) represents the shape of the general profile common to all genes in the group (see S1 Fig). To estimate 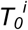, for each gene we compute the correlation between the model *Y^i^*(*t*) and the measured expression timecourse for a range of time-shifts 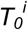, and use the point of highest correlation as the best estimate of the individual time-shift (Fig 1). The advantage of this method is that the expression time for each gene is inferred from many data points, which greatly improves the accuracy of estimation and reduces sensitivity to noise [11,15]. On the other hand, the method can only be used for genes with similar shapes of expression profile, thus potentially restricting the application to a subset of the genes in question.

**Fig 1.**
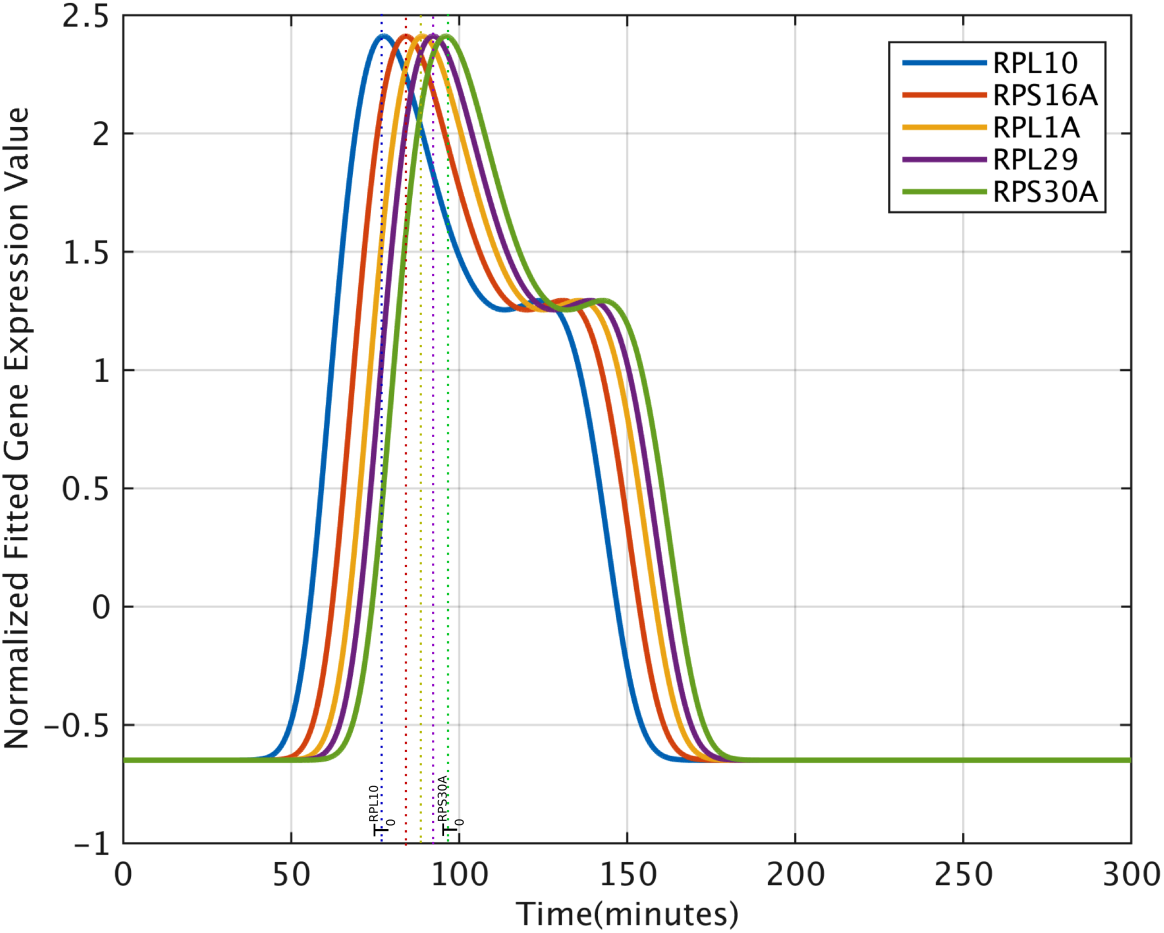
The logic of the model-based timing of r-protein gene expression. Representative fitted curves of ribosomal protein gene expression on the YMC microarray dataset showing early to late peak times. Expression times for individual genes are inferred from position of the best-fit curve.

### The precise timeline of expression of ribosomal protein genes

To infer the timeline of expression of ribosomal protein genes, we use their expression profiles during YMC. During YMC, most of the ribosomal genes are strongly regulated and share a specific shape of temporal expression profile. Our primary dataset is the microarray-based study of [11] that contains more time-points than the more recent RNA-Seq dataset [16]. Other datasets (including cell-cycle synchronized cultures) were not considered because in those systems, the ribosomal genes either are not strongly regulated and correlated, or their regulation is not specific (a high correlation is noted between the expression profiles of r-protein genes and other unrelated genes, (See S1 File and S2 Fig). Therefore, they do not have this obvious consensus pattern fit to our model.

187 genes were selected from 320 candidate genes annotated with GO term “ribosome” (GO:0005840) according to the correlation in YMC expression profiles between these candidate genes and the 114 RPL/RPS genes (see Methods, S1 Table and S3 Fig). Some genes associated with ribosome biogenesis are expressed during YMC in a short burst, which is not strongly correlated with broad peak characteristics of the r-proteins. The YMC temporal profiles of these genes do not fit our model, and are not suitable for timing using our approach and thus these genes were excluded from our following analysis. Therefore, the relative timing of ribosomal gene expression is reliable only for 187 of the genes associated with ribosome biogenesis. Most of these genes are associated with the mature ribosome.

For each of the 187 ribosomal transcripts defined above, we estimated the time-shift *T*_0_^(g)^ at which the model best fits the measurement. This quantity is readily interpretable as the time of maximum expression for each gene. The estimated timeline of r-protein gene expression spans approximately 25 minutes. The median estimated accuracy of timing of a gene is 1.3 minutes, which allows resolving the differences between individual genes (fig 2). The timing results provided in S1 Table. The timing results are more precise for genes with higher correlation between the model (the consensus profile) and the measured expression profile (S4 Fig).

**Fig 2.**
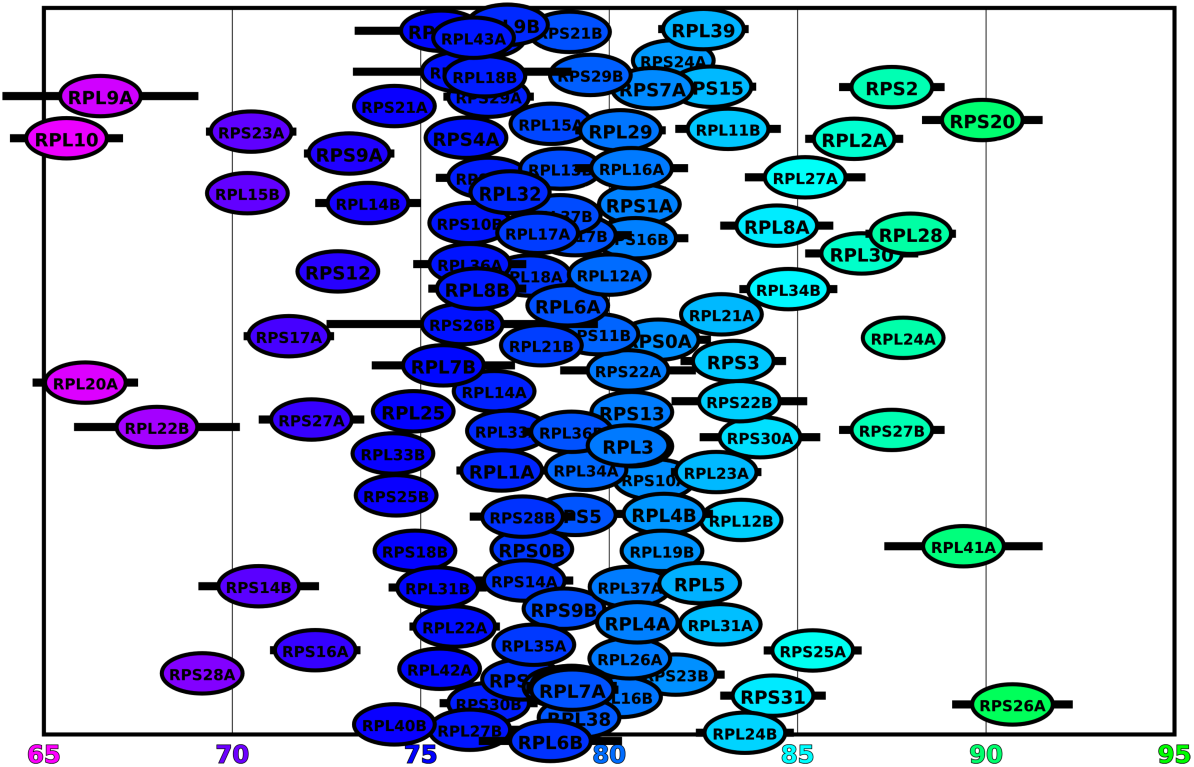
Timeline of expression peaks of r-protein genes during YMC.

By comparing the expression times of the r-proteins in the mature ribosome structure with all proteins involved in ribosome biogenesis, we found that the r-proteins, incorporated into small (RPLs) and large subunits (RPLs), are typically expressed significantly later (*P*=1.02e-8, rank-sum test, one-tailed) than other timed proteins, most of which are not incorporated into the ribosome, see Fig. 3A (this set also contains stalk proteins). GO term enrichment analysis show that most of the proteins not incorporated into LSU or SSU are enriched in ribosome rRNA processing, assembly, and localization, gene expression and translational initiation (see S2 Table). This is consistent with the general role of the structural r-proteins in the last step of assembly [16], while not incorporated proteins generally function in early stage ribosome biogenesis. We further compared the expression times of ribosomal small subunit protein genes (RPSs) and large subunit protein genes (RPLs), The average expression times of small and large subunit transcripts are 78.8 and 79.4 min, respectively and show no significant difference (Fig 3B, *P* = 0.81, Wilcoxon rank sum test, one tailed, *P* = 0.26; t-test). The results is consistent with the fact that the biogenesis of the small and large subunits of the ribosome is simultaneous [36]. Also, when we compare the expression of the RPLs and genes encoding components of ribosomal stalk (RPP0, RPPA1, RPPA2, RPPB1, and RPPB2), we did not observe significant difference of the two group genes in expression times (*P*=057, Wilcoxon rank sum test, one tailed; *P*=0.73, *t-test*). This result does not indicate either that the stalk needs to be completed before LSU or vice versa, and is consistent with independent assembly of the ribosomal stalk and the ribosomal large subunit, although the stalk is finally attached to the ribosomal large subunit in the mature ribosome [37].

**Fig 3.**
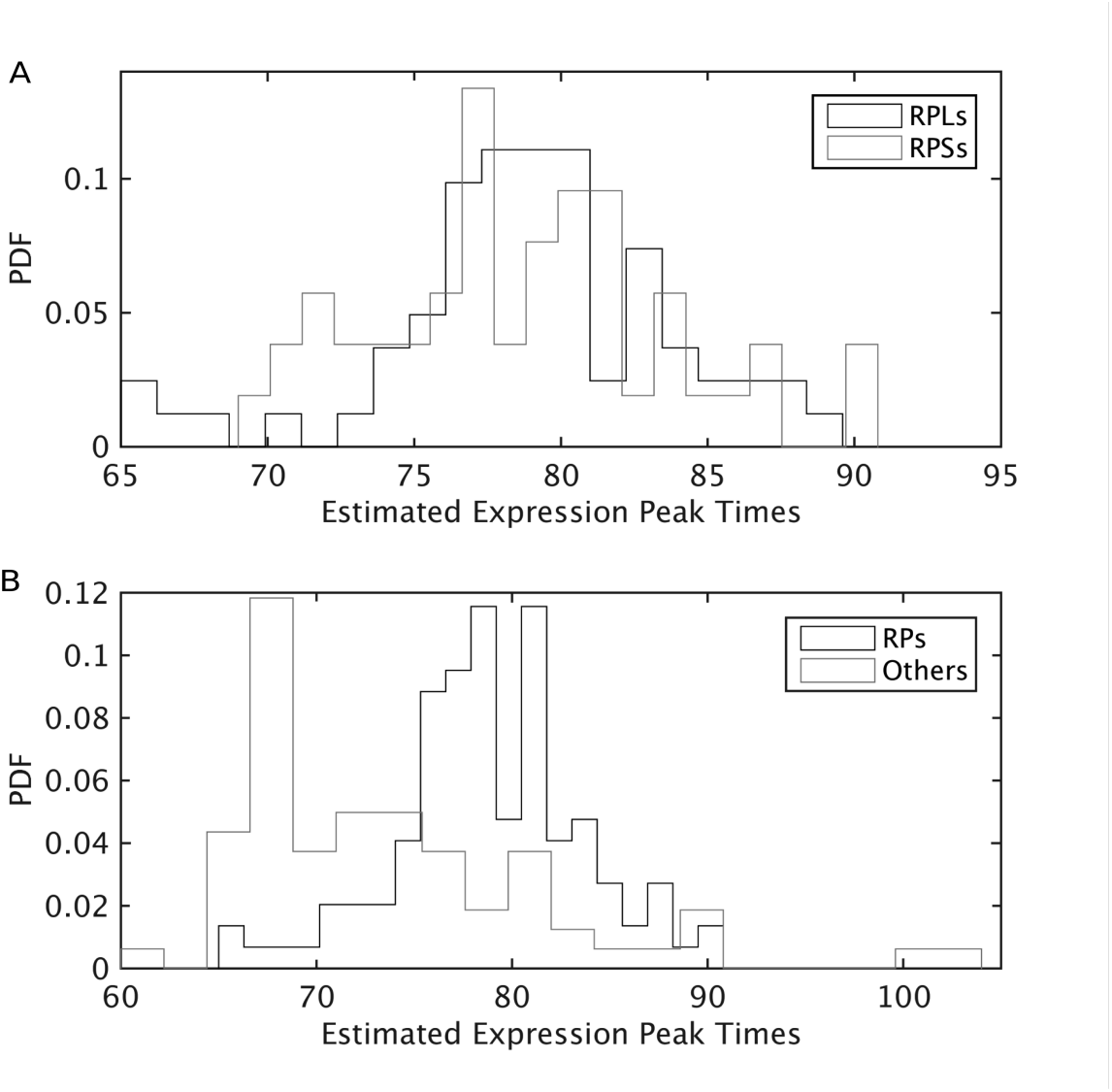
Comparison between expression times of small and large subunit r-proteins (A) and between expression times of r-protein genes and non r-protein genes (B). Panel A: No significant difference between expression times between two groups, i.e., small and large subunit r-protein genes. Panel B: Significant differences between expression times of genes encoding r-proteins integrated into mature ribosome and other r-protein associated genes not integrated into mature ribosome.

### Relation between expression timeline and the causal dependency between ribosomal proteins

Causal dependencies between recruitment of r-proteins during biogenesis of the mature ribosome have been reported in the literature. Depletion of the yeast ribosomal proteins RPL16 (L16) or RPS14B (rp59), disrupts ribosome assembly, and several ribosomal proteins are rapidly degraded in the absence of RPL16 or RPS14B [38]. These reports suggest that RPL16 and RPS14B may be used in ribosome biogenesis before the other proteins. Our expression timing results point to expression of RPS14B before its target genes. All 13 genes dependent on RPS14B (YJL191W) are expressed after RPS14B with *P*-value <1.0e-16 (binomial test) (see Table 1). It is therefore possible that the timeline of expression of these genes agrees with their timeline of incorporation in the ribosome. The same genes were also reported as dependent on RPL16, however 12 of them are localized in SSU, while RPL16 is a component of the LSU, so a direct causation is less likely since SSU and LSU are thought to be independent in biogenesis [36]. Indeed, only two pairs with consistent dependency relationships were observed (see Table 1B). One interpretation is that association between time lines and dependency relationship may be limited to dependencies between pairs of proteins that have neighboring localizations within the structure, or at least are localized within the same ribosomal subunit (here small ribosomal subunit).

**Table 1.**
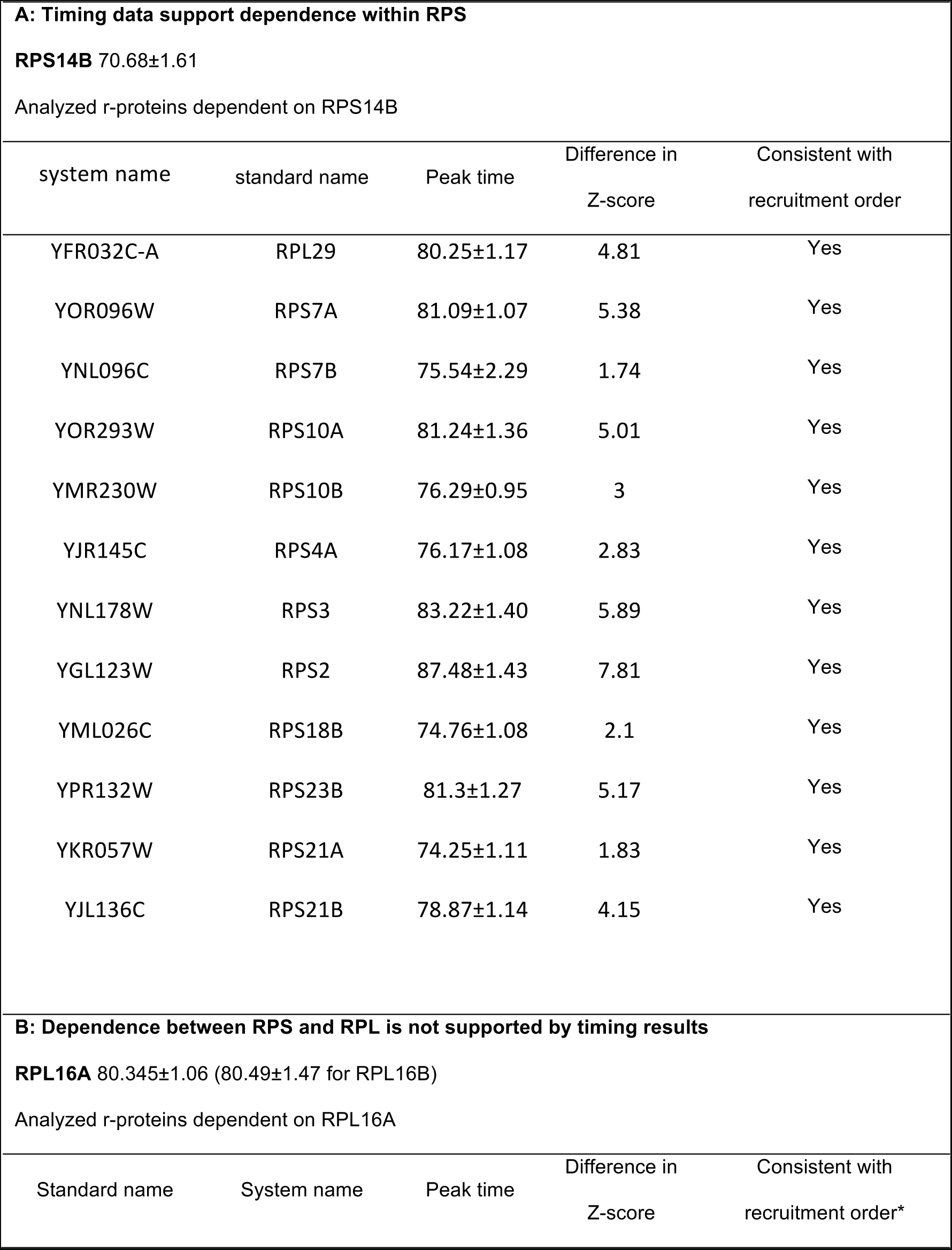
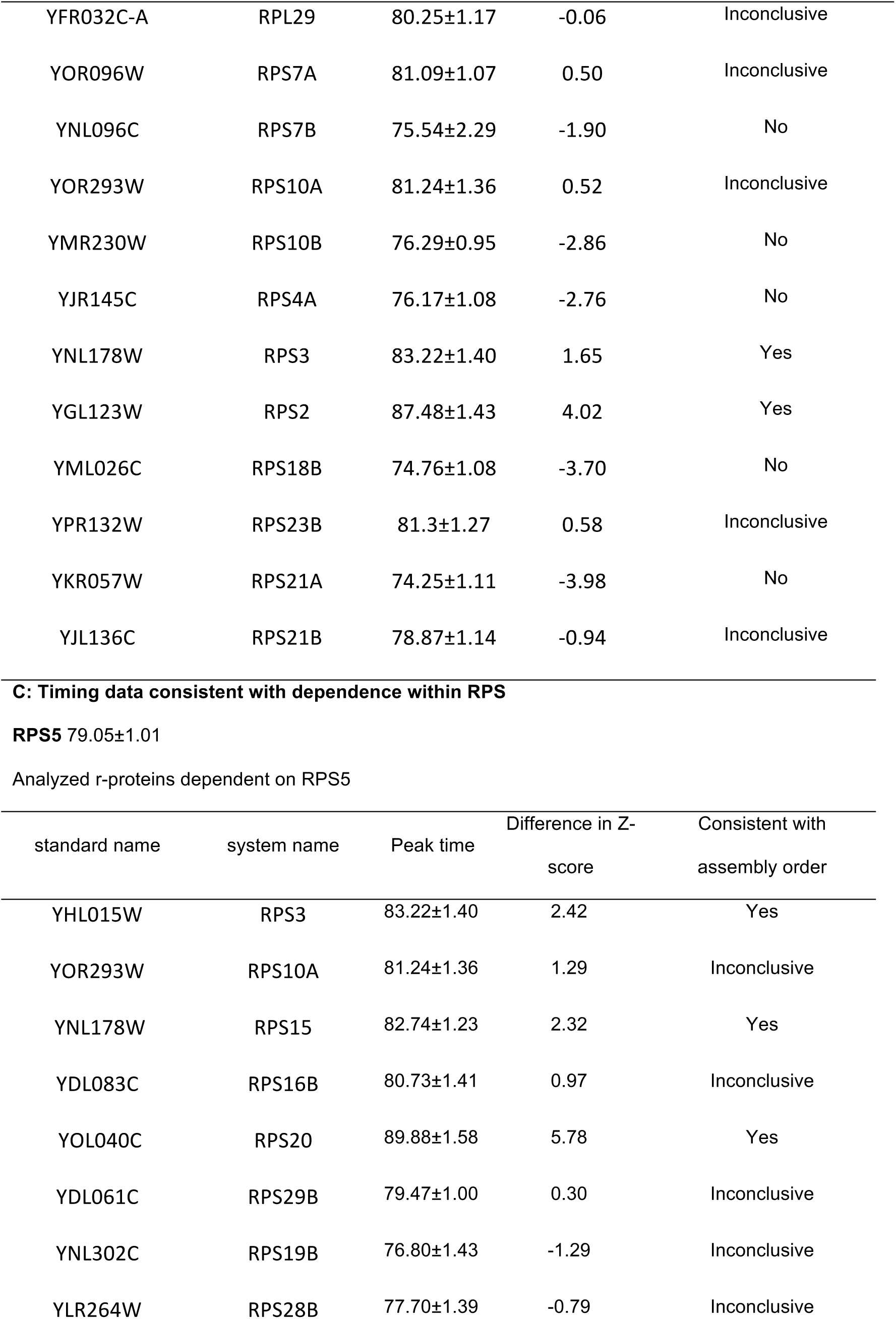
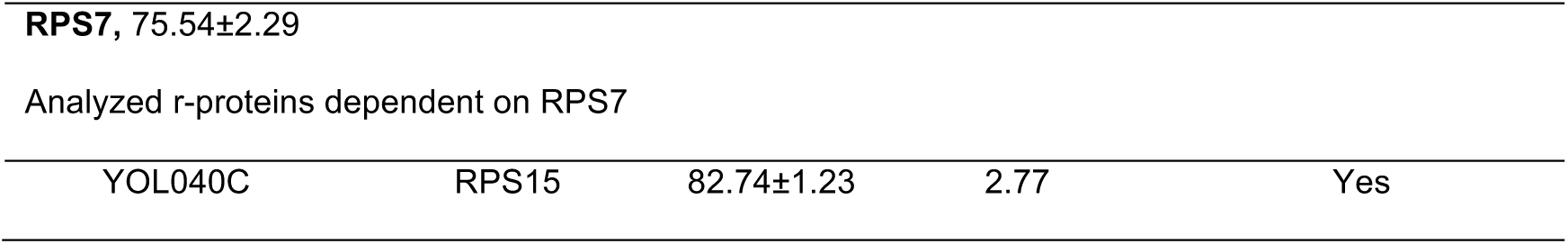
Time of gene expression peaks is significantly associated with the recruitment order as reported in [38] and [20] in ribosome maturation. The order of five pairs of r-proteins of small ribosome subunit in assembly from [20] were analyzed. 12 r-proteins were analyzed that dependent on RPS14 and RPL16 from [38]. R-proteins dependent on RPS7 and RPS5 [20,21] were also analyzed.

| system name | standard name | Peak time | Difference in<br>Z-score | Consistent with<br>recruitment order |
| --- | --- | --- | --- | --- |
| YFR032C-A | RPL29 | 80.25±1.17 | 4.81 | Yes |
| YOR096W | RPS7A | 81.09±1.07 | 5.38 | Yes |
| YNL096C | RPS7B | 75.54±2.29 | 1.74 | Yes |
| YOR293W | RPS10A | 81.24±1.36 | 5.01 | Yes |
| YMR230W | RPS10B | 76.29±0.95 | 3 | Yes |
| YJR145C | RPS4A | 76.17±1.08 | 2.83 | Yes |
| YNL178W | RPS3 | 83.22±1.40 | 5.89 | Yes |
| YGL123W | RPS2 | 87.48±1.43 | 7.81 | Yes |
| YML026C | RPS18B | 74.76±1.08 | 2.1 | Yes |
| YPR132W | RPS23B | 81.3±1.27 | 5.17 | Yes |
| YKR057W | RPS21A | 74.25±1.11 | 1.83 | Yes |
| YJL136C | RPS21B | 78.87±1.14 | 4.15 | Yes |

| Standard name | System name | Peak time | Difference in<br>Z-score | Consistent with<br>recruitment order* |
| --- | --- | --- | --- | --- |
| YFR032C-A | RPL29 | 80.25±1.17 | -0.06 | Inconclusive |
| YOR096W | RPS7A | 81.09±1.07 | 0.50 | Inconclusive |
| YNL096C | RPS7B | 75.54±2.29 | -1.90 | No |
| YOR293W | RPS10A | 81.24±1.36 | 0.52 | Inconclusive |
| YMR230W | RPS10B | 76.29±0.95 | -2.86 | No |
| YJR145C | RPS4A | 76.17±1.08 | -2.76 | No |
| YNL178W | RPS3 | 83.22±1.40 | 1.65 | Yes |
| YGL123W | RPS2 | 87.48±1.43 | 4.02 | Yes |
| YML026C | RPS18B | 74.76±1.08 | -3.70 | No |
| YPR132W | RPS23B | 81.3±1.27 | 0.58 | Inconclusive |
| YKR057W | RPS21A | 74.25±1.11 | -3.98 | No |
| YJL136C | RPS21B | 78.87±1.14 | -0.94 | Inconclusive |

| standard name | system name | Peak time | Difference in Z-score | Consistent with assembly order |
| --- | --- | --- | --- | --- |
| YHL015W | RPS3 | 83.22±1.40 | 2.42 | Yes |
| YOR293W | RPS10A | 81.24±1.36 | 1.29 | Inconclusive |
| YNL178W | RPS15 | 82.74±1.23 | 2.32 | Yes |
| YDL083C | RPS16B | 80.73±1.41 | 0.97 | Inconclusive |
| YOL040C | RPS20 | 89.88±1.58 | 5.78 | Yes |
| YDL061C | RPS29B | 79.47±1.00 | 0.30 | Inconclusive |
| YNL302C | RPS19B | 76.80±1.43 | -1.29 | Inconclusive |
| YLR264W | RPS28B | 77.70±1.39 | -0.79 | Inconclusive |
| **RPS7**, 75.54±2.29 |  |  |  |  |
| Analyzed r-proteins dependent on RPS7 |  |  |  |  |
| YOL040C | RPS15 | 82.74±1.23 | 2.77 | Yes |

In yeast, RPS3, RPS10, RPS15, RPS16, RPS19, RPS20, RPS28, and RPS29 depend on RPS5 in small subunit head structure assembly [20,21], and RPS15 depends on RPS7. According to the justin-time expression paradigm, we expect RPS5 and RPS7 expressed no later than the dependent proteins. Indeed, out of those nine causal dependencies, no pair showed significantly earlier expression for the dependent r-proteins, while in four cases the downstream protein was expressed significantly later than the upstream one. A likelihood calculation yields L=0.0046 for the actual timing, while a reversed timing would provide L= 1.02e-28. Therefore the odds of concordance with timing are 4.46e25 to 1 (Table 1).

### Time of gene expression is correlated with relative position within the structure

Our analysis suggests that expression timing is relevant to the functions of the r-proteins in global ribosome assembly and biogenesis at least for proteins within the same ribosomal subunit. An association therefore likely exists between the expression times and distributions of r-proteins in the mature ribosome. This association could be relevant to functions and sequential association of the r-proteins in the ribosome assembly. To test this, we measured the correlation between difference in mass center distance and difference in time of expression for pairs of ribosomal proteins in the same neighborhood. Our rationale is that the assembly order is determined by spatial constraints. A significant correlation may indicate that at least some of ribosomal proteins, if not all of them, follow the just-in-time expression to facilitate the assembly process.

Specifically, we obtained the atom coordinates of the 53 timed ribosomal proteins within the mature ribosome from the x-ray crystal structure of yeast ribosome [1]. We defined the radial coordinate of each protein as the distance of the mass center of the protein to the mass center of the whole ribosome structure. We considered all pairs of timed ribosomal proteins with an upper limit on the angle θ between the axes connecting the two mass centers of each protein and the whole ribosome center.

For each pair, we noted the difference in radial coordinates (*dr*) and in expression times (*dt*). To characterize the dependence between radial coordinate and expression time, we computed the correlation between these two values over all pairs of neighboring ribosomal proteins. We found that the magnitude and significance of correlation depends on the maximum angle *θ_max_*. For small angles there are too few pairs within a narrow cone for a statistically significant relation, for the widest angles the large distances between proteins influence the statistic result. We tested a range of practical apertures from θ_*max*_=20° to θ_*max*_=90° (using the center of mass of the respective subunit in the calculation). For small and large subunits, the Pearson correlation coefficient are most significant (0.36 and 0.32 with *P*=8.69e-6 and *P*=0.0039, respectively) at angle thresholds of 54 and 42 degrees (Fig 4). The result is similar for Kendall τ correlation coefficient (Kcc) and Spearman correlation coefficient (Scc). For the small subunit, the significant angle threshold of Pcc, Kcc and Scc are the same and range from 36 to 90 degrees (Table 2 and S3 Table). For the large subunit, the Pcc is significant for angles from 37 to 48 degrees; Kcc and Scc are significant between 41 and 44 degrees, as shown in Fig 4 and Table 2. Our results show that the correlation is significant for both small and large ribosome subunits, with more significance for SSU. The expression times of r-proteins in the interface and the solvent-exposed surface of the small and large subunits of the ribosome (LSU) are respectively color-annotated with heatmap in the ribosome structure, and shown in Fig 5(A, B) and Fig 5(C, D), where the early expressed r-proteins are denoted by purple, while the late ones are color-coded green.

**Fig 4.**
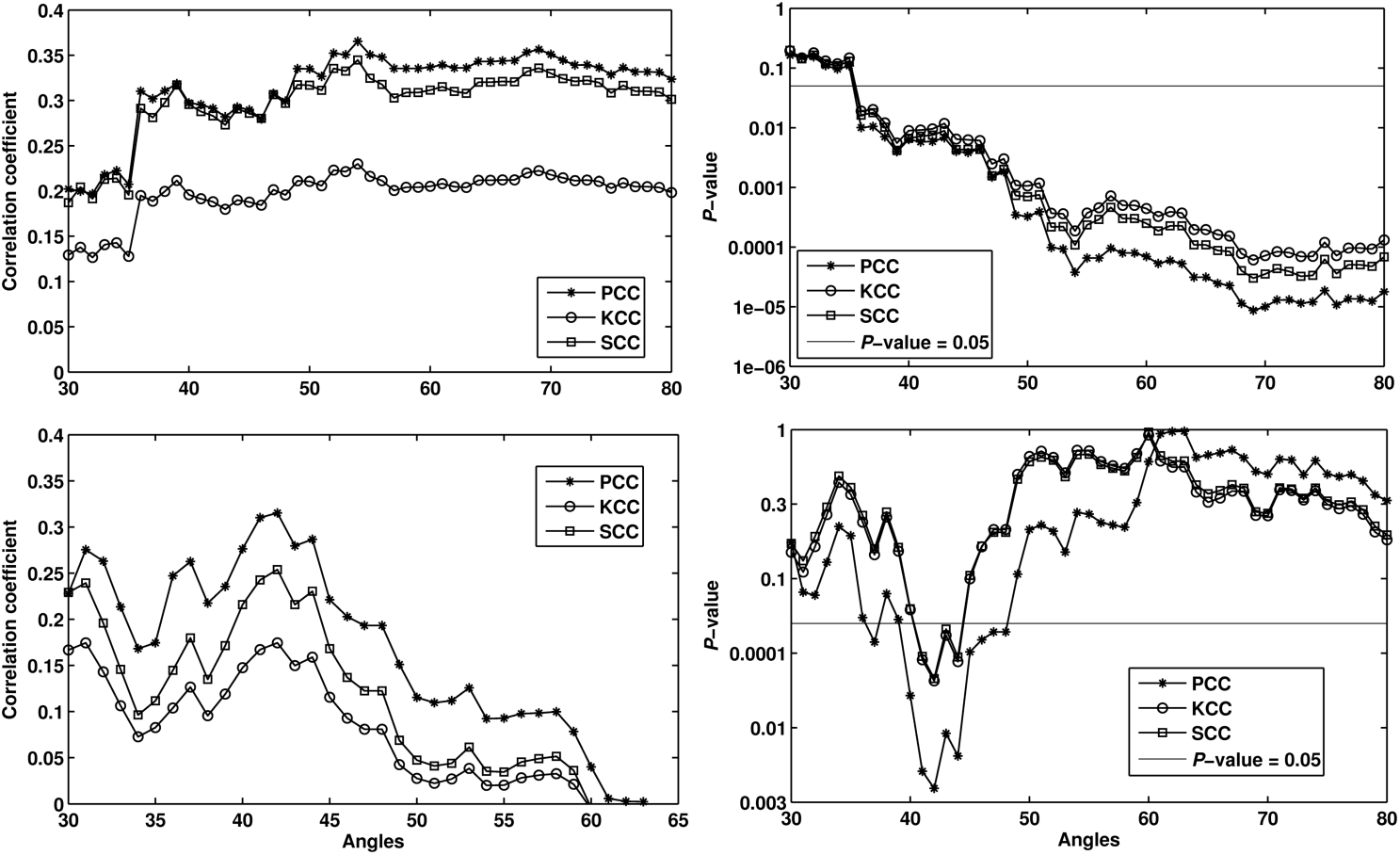
Plot of Pearson, Kendal and Spearman correlation coefficients versus angle thresholds between the difference in mass center distance (*dr*) of each pairs of proteins in ribosome small (top panel) and large (lower panel) subunit structure under a specific angle threshold and the difference of protein transcription peak timing (*dt*). PCC, Pearson correlation coefficient; KCC, Kendal correlation coefficient; SCC, Spearman correlation coefficient.

**Table 2.**
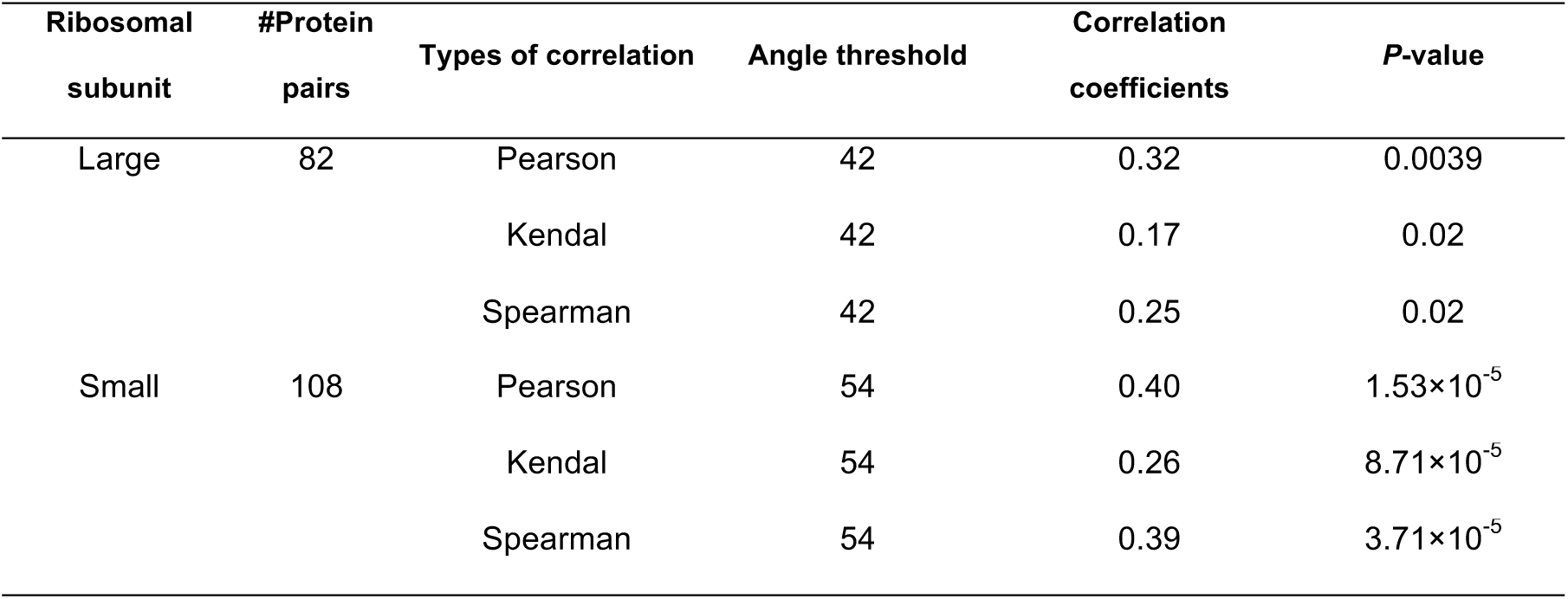
Correlation coefficients between protein mass center and timing at the most significant angle threshold between observed protein pairs. Information on the examined angle thresholds can be found in S3 Table.

**Fig 5.**
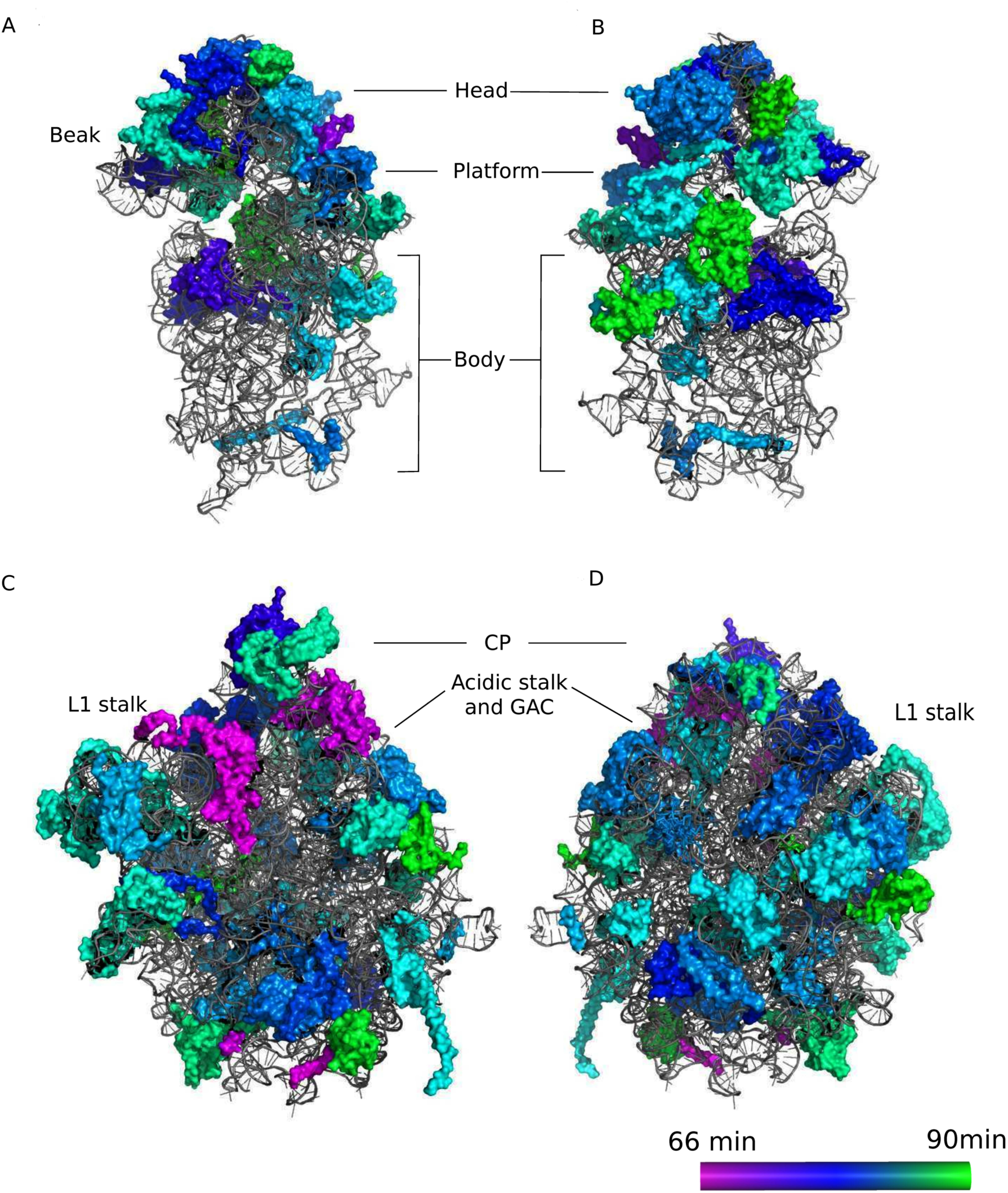
Timeline of transcription peak time of each gene in the mature ribosome structure (PDB code: 3O30 and 3O5H). (A,B) The interface and the solvent-exposed surface of small subunit ribosome, respectively; (C,D) the interface and the solvent-exposed surface of large subunit ribosome, respectively. The expression time of ribosomal proteins is color-coded from early (purple) to late (green). Abbreviations: CP, central protuberance; GAC, GTPase-activation center.

### Lack of significant correlation of the timing with linear position along the rRNA

Reports suggest that early stages of ribosome assembly are co-transcriptional with rRNA [39], which may indicate that r-proteins associated with 5’ RNA may be incorporated earlier than those associated with 3’ RNA, as observed in bacteria [40]. Our result shows no significant correlation of the timing with linear position along the rRNA (see supplementary materials and S5 Fig for more details) and thus does not indicate that co-transcriptionally recruited proteins are expressed in the same general order in which they are used. Note however, that the ribosomal genes with reliable timing are mostly those associated with the mature ribosome, and many of them may not be expressed co-transcriptionally with the rRNAs. Conversely, many pre-ribosomal proteins that are presumably co-transcriptional have their mRNA expressed in a short burst and therefore their expression time during YMC could not be assessed using our model.

### Regulatory sequence elements

The presented results are consistent with the proposition that the time of ribosomal gene expression may be important for ribosome biogenesis. It is therefore expected that a regulatory mechanism may exist that controls the time of expression. It has been argued that in yeast, the expression profiles of genes are highly determined by the regulatory elements in the promoters [8]. One can therefore expect that the expression times may also depend on the regulatory motifs. We characterized the regulatory elements in the 5’ regions adjacent to the 187 ribosomal genes using the MEME motif identification software [41]. The significant motifs present in 30 or more of the 187 genes are summarized in Table 3. Most of the enriched motifs were previously reported in r-protein gene expression regulation, such as the RAP1/SFP, A-rich, DAT1 motif (nonalternating oligo(A).oligo(T) tracts, i.e., A.T tracts), NHP6A motif (AT rich high-mobility motif, NHP6A is homologous to human HMG1 and HMG2), and CG rich motifs (S4 Table).

**Table 3.**
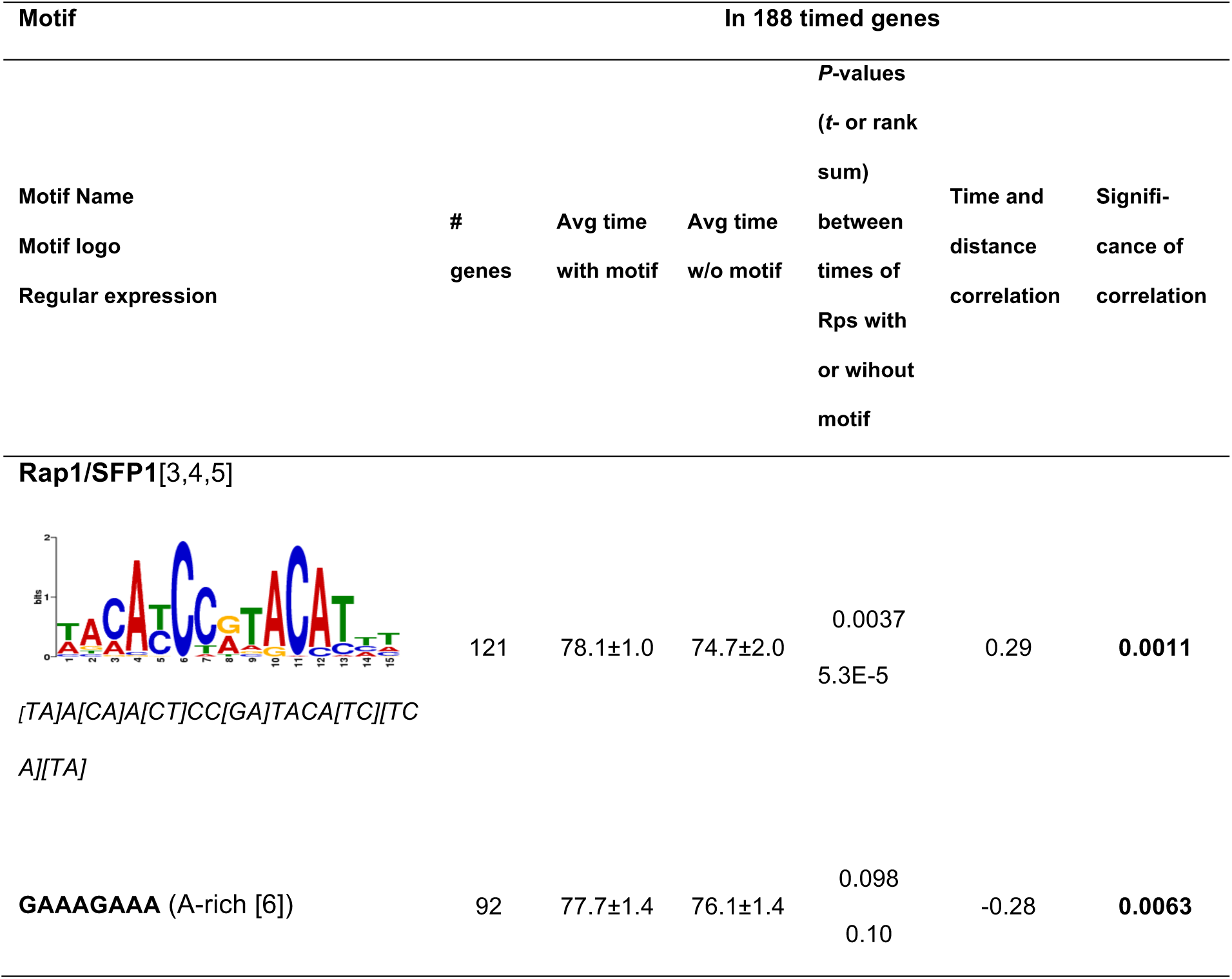
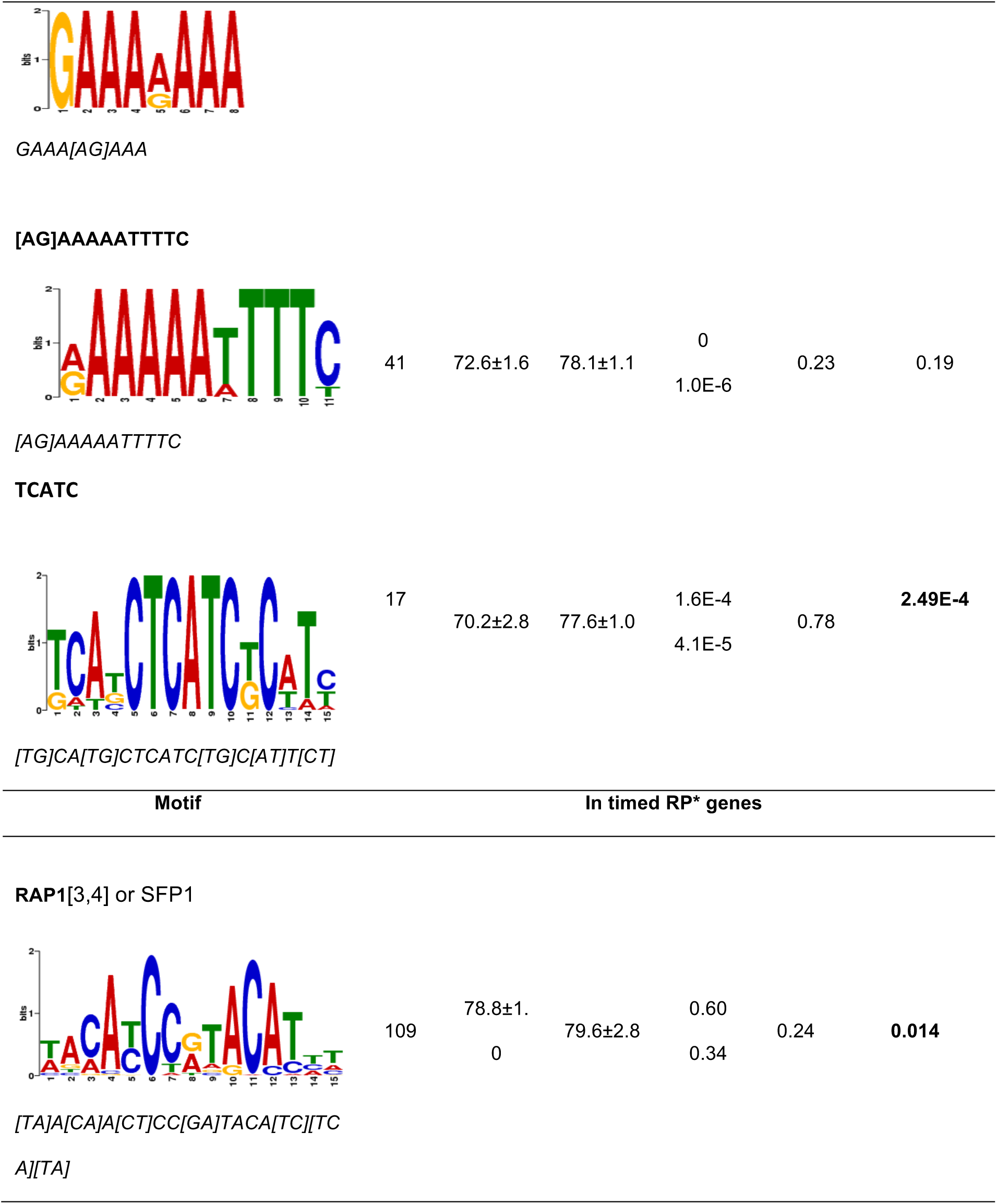
The significantly correlated regulatory elements potentially involved in regulating the times of ribosomal gene expression. Top panel: all 187 timed genes; bottom panel: ribosomal genes with ‘RPL’ or ‘RPS’ as the first three characters of their common gene names. Column 1, motif name, logo and the regular expression defining the motif; column 2, number of genes containing the motif; column 3 and 4 are average time of expression for genes with and without motif, respectively; column 5, *P*-value of the *t*-test (first line) and Wilcoxon rank sum test (second line) on the difference in expression time between genes with and without motif; column 6, Pearson correlation coefficient between expression times and distances from motif to coding sequence; column 7, significance of the correlation between motif location and expression time. For all the enriched regulatory elements in the promoters of 187 ribosome related proteins, see S4 Table.

For every motif, we compared the timing of genes with the motif, and genes without the motif, using both *t*-test and Wilcoxon rank sum test. For each motif, we also calculated the correlation between time of expression and distance between motif in 5’UTR and start of the coding sequence.

The well-known RAP1 motif that regulates r-protein gene expression is found in the 5’UTR of most (precisely 109 out of 187) of the timed genes coding for ribosomal and ribosome-associated proteins. Genes with the RAP1 motif were expressed significantly earlier than genes without this element (*P* = 5.3E-5, Wilcoxon test). Moreover, a significant Pearson correlation coefficient (*r*) was observed between the expression times and the distances of the RAP1 motif from the translational start site in the 5’UTR (*r* = 0.29, p-value=0.0011). The Pearson correlation is still significant (*P* =0.01) when only the genes coding for proteins previously annotated as structural components of the ribosome are taken into account (Table 3). Another motif potentially significant for timing of expression is the [AG]AAAAATTTTC element, corresponding to the DAT1 AT tract motif reported in [9]. The genes with the [AG]AAAAATTTTC motif were typically expressed earlier than genes without this element, this difference is very significant in the entire set of 187 timed transcripts (*P* = 1.0E-6, Wilcoxon rank sum test)(Table 3). In addition to the positive correlation, a negative correlation was observed between the motif locations and expression peak times for the GAAAGAAA (A-rich [6]) motif (2^nd^ row in Table 3). While these observed correlations do not prove that the motifs are directly involved in controlling the timeline of expression, our results are consistent with the hypothesis that the timing of ribosomal genes is functional at least in certain conditions and the significant motifs identified here are involved in the process.

Specifically, the correlation between expression time and distance from motif to coding sequence may suggest that a mechanism may be involved that translates the distance to expression time. Alternatively, delaying expression of a ribosomal gene may be caused by the presence of additional elements between the RAP1 motif and the 5’ end of the coding sequence, which will have the side effect of increasing the distance between RAP1 and the coding sequence. In either case, identifying the mechanism and its underlying biophysical details will require further experimental studies.

### Reproducibility of the estimated expression times for the ribosomal protein genes

To validate the robustness and reproducibility of our analysis, we also inferred expression times for the recently published expression profiles in YMC examined in a RNA-seq experiment [16]. The profiles are strongly correlated with the model derived from microarray data (average Pcc=0.785), so the model could be applied without modifications. The peak times inferred from the two independent datasets are highly correlated (Pearson coefficient r = 0.62, p-value = 4.2e-20). After removing one outlier, the correlation reached 0.72 with p-value = 1.42e-28, see Fig 6 (the outlier is YPP1/YGR198W – a protein identified as possibly interacting with the ribosome based on one copurification study [42], but its primary function is associated with plasma membrane and endocytosis [43,44]),. Peak times inferred based on RNA-Seq are provided in supplementary S5 Table.

**Fig 6.**
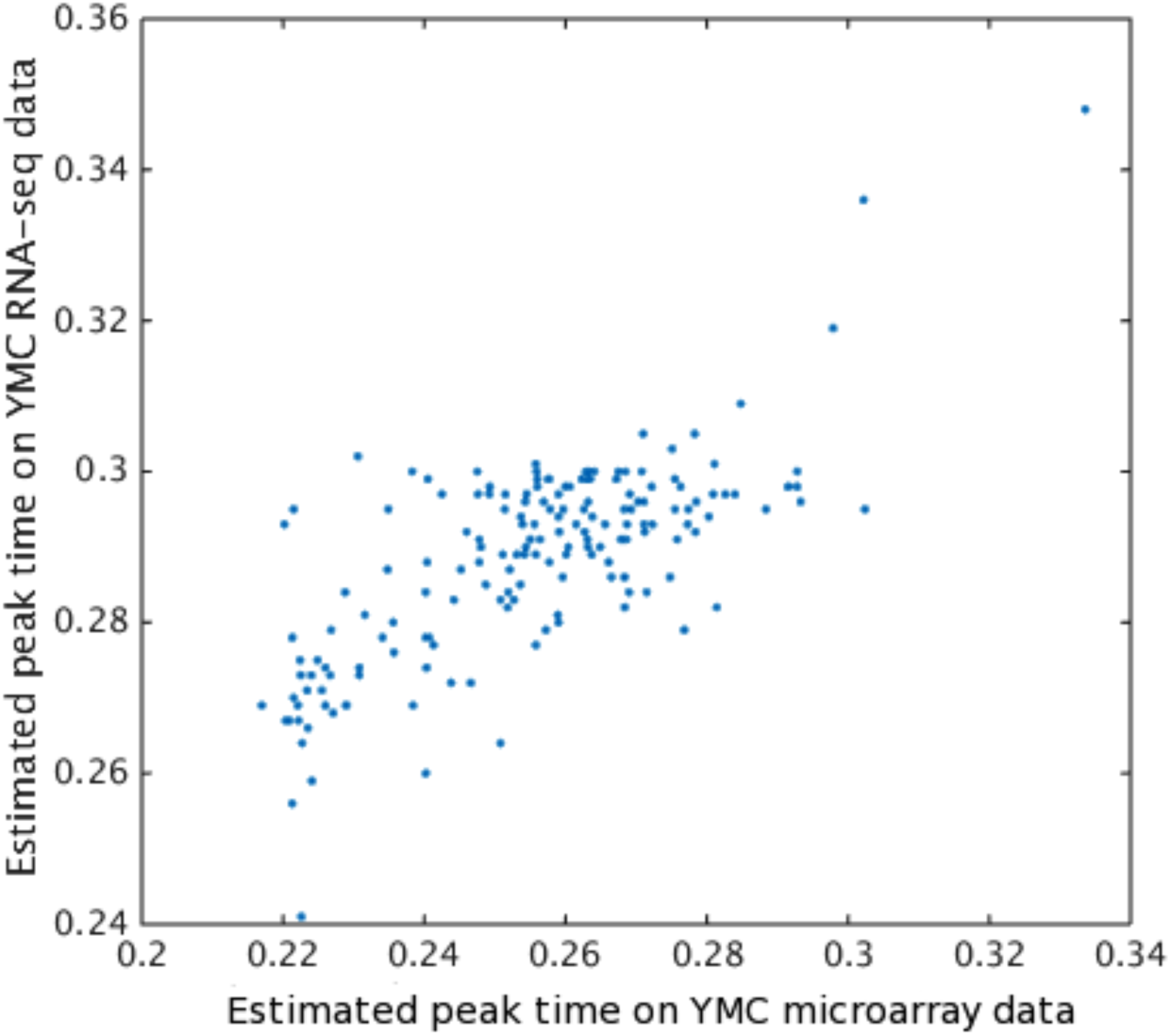
Reproducible estimated expression times of r-protein and associated genes as demonstrated by high correlation coefficients.

This result validates the reproducibility of the result and constitutes evidence supporting possible biological significance of the order of expression of ribosomal genes.

## Discussion

Model-based analysis of biological systems often allows discovering features that are not apparent in the raw data. Here, we applied a new model-based timing algorithm to the expression profiles of ribosomal genes during Yeast Metabolic Cycle. As a result, we obtained a precise timeline of expression of 187 genes coding for proteins associated with ribosome or ribosome biogenesis. While the temporal profiles are similar, these genes are not expressed simultaneously, but rather follow a specific order that is consistent between two experiments. The inferred timeline of expression of ribosomal genes spans approximately 25 minutes, which is longer than duration of assembly previously observed in in-vitro experiments. A possible explanation of the extended timeline is that during YMC the cells may be finely programmed for preparing for the subsequent steps, and the longer timeline may facilitate temporal compartmentalization of the subsequent steps of assembly. Such compartmentalization, or temporal separation of sequential steps in ribosome biogenesis, may play a part in facilitating the fidelity and integrity of ribosome biogenesis.

Through association study of the relative expression time of neighboring r-proteins and their relative distances to the centroid of the ribosome, we found that for pairs of ribosomal proteins from the same region of the ribosome, the protein localized closer to the center of mass of the entire complex tends to be expressed earlier than its neighbors that are more distant from the center of mass of the entire complex. A possible explanation of this correlation is that the ordered timeline of this regulation of neighboring r-proteins may be fine-tuned to facilitate assembly by overcoming spatial constraints during ribosome biogenesis. This is reminiscent of the principle that the index of amino acid depth in a protein complex and its buried surface can be used to identify amino acids that are most crucial for in protein complex formation [45,46]. This hypothesis may extend to the protein level and protein neighbor pairs in ribosome assembly, where the r-protein with mass center closer to the center of the ribosome can have greater contribution to the stability of the entire structure and thus it may be beneficial if it is expressed earlier. Our results may reflect a correlation that may exist also between relative positions of proteins and their order of assembly.

Moreover, although many exceptions exist (e.g. early transcription of RPL10, RPL40 and RPL43), in our cases, the order of expression is consistent with the order of incorporation of r-proteins. This correlation is highly significant especially for the SSU r-proteins. Our observation that the correlation between the mass centers and expression times is more significant for SSU r-proteins than LSU r-proteins when constrained with the neighborhood, may reflect the different complexity of the SSU and LSU biogenesis [47]. While it is not known if the phenomenon is specific to the metabolically synchronized culture during YMC, it is possible that the described correlations are more general, however they could not be observed in other systems in which the time-dependent regulation of ribosomal genes is less prominent, nonexistent, or not synchronized between the cells in the sample.

Ribosomal proteins can play extra-ribosomal roles in gene expression regulation [48,49] through interacting with proteins from other processes. For example, r-proteins may act as modulators of the transcription factor NF-κB activity in gene expression [50,51,52] in mammalian cell lines. Through direct interaction with MDM2 and subsequent p53 activity increase, specific groups of RPL5, RPL11, RPL23, etc., integrate the inhibition of cell growth with cell cycle arrest [53]. Many other studies also found that specific ribosome protein genes are differentially expressed in different tissues [54], between normal and cancer tissues and metastatic and non-metastatic cell lines [55]. Specialized ribosomes have been reported [56] that translate specific transcripts under specific conditions, such custom ribosomes are assembled using different combinations of r-proteins. Also, the properties of ribosomes within the cytoplasm are different from those associated with the endoplasmic reticulum. It is likely that the transcriptional regulation of the r-proteins, including the temporal order of their expression and assembly, may be instrumental in the finer control of the customized biogenesis process and/or extra-ribosomal functions.

Furthermore, significant correlation between the distance of several motifs to the translational start sites and the estimated timelines suggests possible mechanism regulating the timing of the r-protein expression. These include RAP1 and the TGAAAAATTTT motif, which provide testable hypothesis for further validation.

In summary, we have inferred a high-resolution timeline for the r-proteins. We provided evidence in favor of functional relevance of the differential timing. Finally, we pointed to potential functional involvements of the fine temporal compartmentalization of the r-protein gene expression, and to a possible mechanism responsible for regulating the precisely timed expression.

## Materials and Methods

### Yeast r-protein gene time-course expression profiles

The YMC dataset of [14,16] has high data quality (See supplementary materials and S6 Table) and is highly relevant to the task, and therefore, was finally selected as the primary source of expression data for further analysis. Out of the two YMC timecourses available [14] and [16], we selected the [14] as it contains a larger number of samples than the latter in total.

320 genes annotated with GO term “ribosome” (GO:0005840) are selected as coding for potential ribosome assembly related proteins. Correlations in YMC expression profiles between these candidate genes and the 114 RPL/RPS genes are calculated to assess whether they are related to the ribosome biogenesis. According to the correlation with the mean of expression profiles of the 114 genes whose common names start with ‘RPL’ or ‘RPS’ (see supplementary methods), 187 of the 320 transcripts were selected, which constitute a group with a distinctly high correlation to r-protein genes as shown in S3 Fig. The similarity between temporal profile curves was also a prerequisite for the timing algorithm to produce accurate results. Among the 187 genes, 122 are annotated by the GO term “structural constituent of ribosome” (GO:0003735). Notably, this list did not contain any components of the mitochondrial ribosome, which is consistent with the fact that they are not involved in cytosolic ribosome maturation, and are regulated by separate processes. Therefore, all 187 transcripts were included in the timing analysis.

### Model of common expression profile and precise time estimation

Semi-empirical consensus models are widely used to characterize molecular kinetics in gene expression, biophysics [11,57,58] and development [15]. Here, we proposed a novel near Gaussian polynomial model to achieve a high-resolution timeline for the expression of genes encoding the mature yeast r-proteins. Our approach to timing assumes that the temporal profiles of r-protein expression will generally reflect the rate of ribosome production in time. The temporal profiles of ribosomal gene expression profiles will all show the same general pattern, with different additional time-shifts dependent on the time of peak expression.

Therefore, we first constructed the common temporal profile that corresponds to the output of ribosomes in the experimental culture. We derived the empirical formula describing the common shape by comparing the averaged temporal expression profile of the ribosomal genes to a family of modified approximations to near-Gaussian distributions [59,60]. It was determined by a maximum-likelihood parameterization applied to time-course microarray data. We subsequently used this consensus profile in the timing analysis by comparing it against the expression profiles of all respective r-protein genes with different time-shifts. We adopted the time-shift at which the similarity was the greatest so as to estimate the most likely time of the expression peak of each gene in question.

The YMC dataset includes mRNA measurements from three metabolic cycles. The overall duration is approximately 900 minutes and thus each cycle has a duration of 300 minutes. To simplify the problem and reduce the effects of noise, each of the three cycles was aligned and thus merged into one meta cycle by using the peaks of O_2_ consumption as timestamps, see [11]. Based on inspection of the shapes of expression profiles of the ribosomal genes, the following near-Gaussian distribution function *Y(T)* was used as the closed-form approximation of the expression profiles of 114 r-protein genes in YMC data: 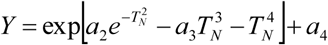, where *T_N_* represents time variable that is scaled by a common factor and shifted to match an individual gene’s delay in expression. *T_N_* depends on the wall-clock time *T* through 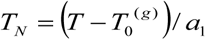, where 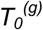 is the gene-specific time shift of expression profile, and *a*_1_, *a*_2_, *a*_3_ and *a*_4_ are global parameters that need to be optimized. With a target function derived from a least-squares fit, we used the conjugate gradients [61] optimization method to minimize the RMS distance between model and expression profile of each r-protein. The resulting consensus temporal expression profile is:

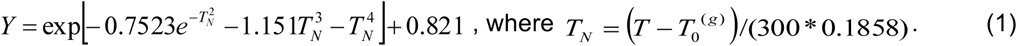

The individual time-shifts of every transcript, 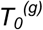, reflect the estimated difference between the expression times of each ribosomal gene in minutes. These time-shifts, computed for the 187 selected gene transcripts, represent the timeline of expression of r-proteins and other proteins in the final stages of ribosome assembly (see Fig 2), and constitute the main result of this study. The computer program for computing the 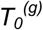 is included in the supplementary materials. The average correlation between the model and the corresponding expression profile is very high and equals 0.78. Our precision estimation result shows that the timing accuracy is higher for profiles strongly correlated with the model, and poor if the profile is not well described by the consensus model (See supplementary materials and S4Fig).

### Association analysis of the times of expression and order of recruitment during ribosome assembly

We used the data from [20,21,38] to validate the biological significance of the times of expression of the r-proteins. When an r-protein was mapped into multiple genes encoding two paralogs, two times of expression will be obtained. We chose one most consistent with the assembly order in that one paralogue of an r-protein can probably displace another one in the process assembly. Therefore, our estimated times of expression can be assessed for consistency with the just-in-time expression.

### Three-dimensional structure of the ribosome

For relating the timing results with the structure of the ribosome, we used the 4.5 ångström resolution crystallographic structure of the fully ratcheted state published by [1]. We collected the atomic coordinates from the Protein Data Bank (PDB - http://www.rcsb.org/pdb/) under accession numbers 3O30 and 3O5H. These structures contain four rRNA chains and 53 defined protein chains: 1 18S rRNA and 20 defined protein chains in the small subunit 3O30 and 3 rRNA (25S, 5S and 5.8S rRNA) and 33 defined protein chains in the large subunit 3O5H. This structure was used to identify relative positions of protein mass centers, i.e., relative distances from the centroids (mass centers) of the ribosomal proteins to the ribosome’s centroid (mass center), as well as contacts between ribosomal proteins and ribosomal RNA.

### UTR sequence analysis

Previous studies indicate that the gene linear structure and transcription factor bindings affect gene expression timing. We collected the sequences of the UTRs (untranslated regions) of 1,000 bp upstream of the 5’ end of the coding region for each of the 187 co-expressed genes. The regulatory elements and their positions with respect to the 5’ end of the coding sequence were identified *de novo* using Meme [41]. The command line input parameter is: *meme -p 24 –mod tcm –recomp –nmotifs 16 –minw 6 -maxw 18*.

For evaluating the significance of the differential expression time between genes with different features in the promoters, we used the Wilcoxon rank sum test throughout the paper. The rationale for this is that we expect that the order of expression (rather than the absolute time) is more important for ribosome assembly, therefore a rank-based test would be the most appropriate.

### Reproducibility of the estimated expression times for the ribosomal protein genes

We also applied our fitted model based on the microarray data of r-protein gene expression profiles directly to the independent RNA-Seq data [16] more densely sampled but with fewer points. The Pearson correlation coefficient between the estimated peak times from RNA-Seq data [16] and microarray data respectively were calculated. A high and significant Pearson correlation coefficient would indicate the reproducibility of our fitted model, and vice versa.

## Acknowledgment

We thank Ben Tu for fruitful discussions, Chuanying Chen and B. Montgomery Pettitt for structural visualization discussion and Heather M, Lander for comments and editing of the manuscript.

## Supporting Information

All relevant data and methods are within the paper and its supplementary materials, as well as on the supporting website: http://moment.utmb.edu/ribosome.

### Supplementary Tables

**S1 Table.** Timing of expression of 187 ribosomal proteins and proteins associated with the mature ribosome.

**S2 Table.** GO Term enrichment analysis and annotation of the proteins not incorporated into the mature ribosome proteins.

**S3 Table.** The Pearson, Kendal and Spearman correlation coefficients versus angle thresholds between the difference in mass center of each pair of proteins (*dr*) under a specific angle threshold and the difference of protein transcription peak timing (*dt*) in whole ribosome structure.

**S4 Table.** The regulatory elements involved in regulation of ribosomal genes. Column 1, motif logo and the regular expression defining the motif; column 2, number of genes containing the motif; column 3 and 4 are averaged times of expression for genes with and without motif, respectively; column 5, *P*-value of *t*-test (top line) and Wilcoxon rank sum test (bottom line) of the difference in expression time between genes with and without motif; column 6, *P*-value of *t*-test and Wilcoxon rank sum test between motif locations of early (earlier than 75 min) and late (later than 75 min) genes; column 7, Pearson correlation coefficient between expression times and distances from motif to coding sequence; Column 8, significance of the correlation between motif location and expression time.

**S5 Table.** RNA-seq based timing of expression of 187 ribosomal proteins and proteins associated with the mature ribosome.

**S6 Table.** Description of the compared 10 datasets. Correlation of expression profiles of ribosomal genes among themselves and with other genes in different datasets.

### Supplementary figures

**S1 Fig.** The consensus meta-profile (in red) fitted to the expression profiles of 187 ribosomal proteins and associated genes in the YMC microarray dataset. The meta-profile is used for the down-stream time shift estimation of each gene’s expression peak by sliding with an increment of one time unit.

**S2 Fig.** Histogram of the correlations of the expression profiles of 320 genes annotated with GO term “ribosome” (GO:0005840) with the averaged expression profile over the selected 114 genes encoding the co-expressed proteins associated with ribosome assembly. The genes with the high correlation as indicated by the red square were selected in this study.

**S3 Fig.** The strong modulation of r-protein gene expression observed in yeast metabolic cycle dataset, compared to other datasets of yeast cell cycles.

**S4 Fig.** The timing error depends on the profile quality, where the high correlation leads to less error. X-axis: correlation (*C*) between model and data. Y-axis: standard deviation of timing in simulated profiles in one unit of cycle length (300 min). Points: 5,100 individual simulated profiles.

**S5 Fig.** Lack of significant correlation with linear position along the rRNA. (a) 25S rRNA (b) 5S rRNA; (c) 5.8S rRNA; (d) 18S rRNA.

